## Supplemental Data for "Neural speech restoration at the cocktail party: Auditory cortex recovers masked speech of both attended and ignored speakers"

### Simulation of TRF cross-contamination

One aspect of the results presented in the main paper concerns estimating TRFs to onsets in the ignored speech source, above and beyond onsets in the acoustic mixture (Figure 3). This deserves special care because these two predictor variables are not orthogonal. Specifically, there is a concern that a true response to the mixture might cause a spurious response to the ignored source. Simulations designed to address this concern were run in order to determine whether such cross-contamination might be seen.

<u>Methods</u>

Simulations were performed using the same stimuli used in the MEG experiment, with the envelopes and onsets downsampled to 100 Hz. Simulated TRFs were manually predetermined, using a combination of Gaussian windows, to resemble experimental TRFs to the mixture and the attended source of envelopes and onsets, respectively (red lines in Figure S1). No TRFs were assigned to the ignored source, because detecting spurious TRFs in the ignored source was the aim of this simulation-based investigation. The simulation encompassed 40 simulated subjects. For each subject, attention alternated between the female and male stimulus for the stimulus pairs 1-4 (order counterbalanced as in the MEG experiment). Each stimulus pair was used only once, i.e., each simulation used 4 minutes of data. The response was simulated by convolving each of the 256 bands of each predictor with the TRF corresponding to the predictor, summing all responses, and adding pink noise based on the Voss-McCartney algorithm, with a signal to noise ratio (SNR) of 1:10 (i.e., -10 dB SNR). TRFs were then reconstructed with the same algorithm as in the main experiment, and the same predictors binned into 8 frequency bins.

<u>Results</u>

Figure S1 displays the simulated TRFs (thick red lines) overlaid on the STRFs estimated in the simulation (blue/yellow lines with standard error). Onset TRFs exhibit no notable spurious response to the ignored speaker, suggesting that leakage from onset responses to the mixture is unlikely to be the source of the results obtained in the MEG study.

The results from the experimental MEG responses, discussed in the main paper, indicated a systematic difference in response function latencies between the onset mixture representation and the individual streams. Thus, a secondary concern arose as to whether peak latencies to those representations systematically affect each other. A second set of simulations addressed this (Figure S2). The simulations were identical to the previous ones, except that the early onset peaks were somewhat narrower, and the latency of the attended stream response was systematically shifted to be earlier than, simultaneous with or later than the mixture response (indicated by the vertical dotted lines). Results indicate that each peak might be slightly mislocalized towards the latency of the other peak, i.e., if any bias were present, it would have worked against the results observed in the experimental responses.

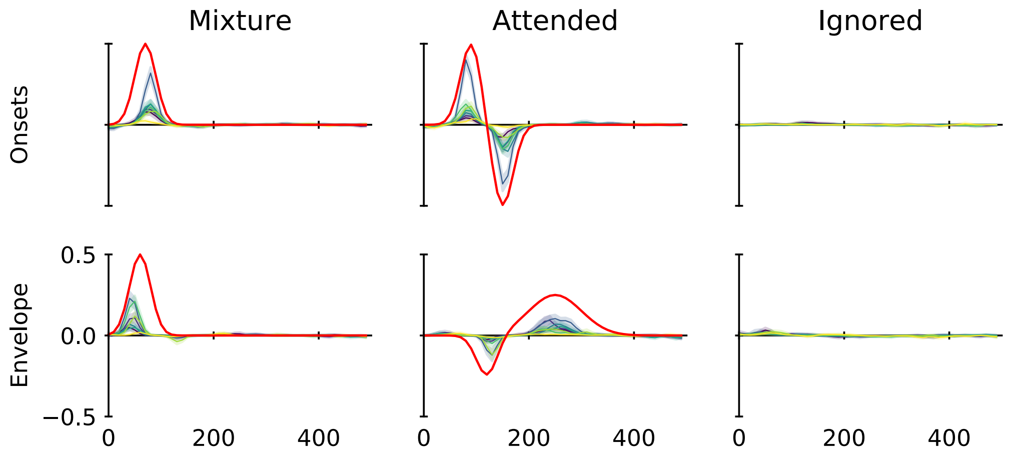


Figure S1. **Simulation without response to ignored speech.** Simulation with conditions selected to approximate results obtained in the MEG study shown in Figure 3. Solid red lines indicate the simulated TRFs; yellow/blue lines show the STRFs reconstructed with methods analogous to the MEG analysis. Note the absence of spurious responses to ignored onsets, indicating that the algorithm used here is robust against false positives regarding ignored onsets. Shading indicates within-subject standard error of the mean. Simulated and estimated TRFs are at different scale.

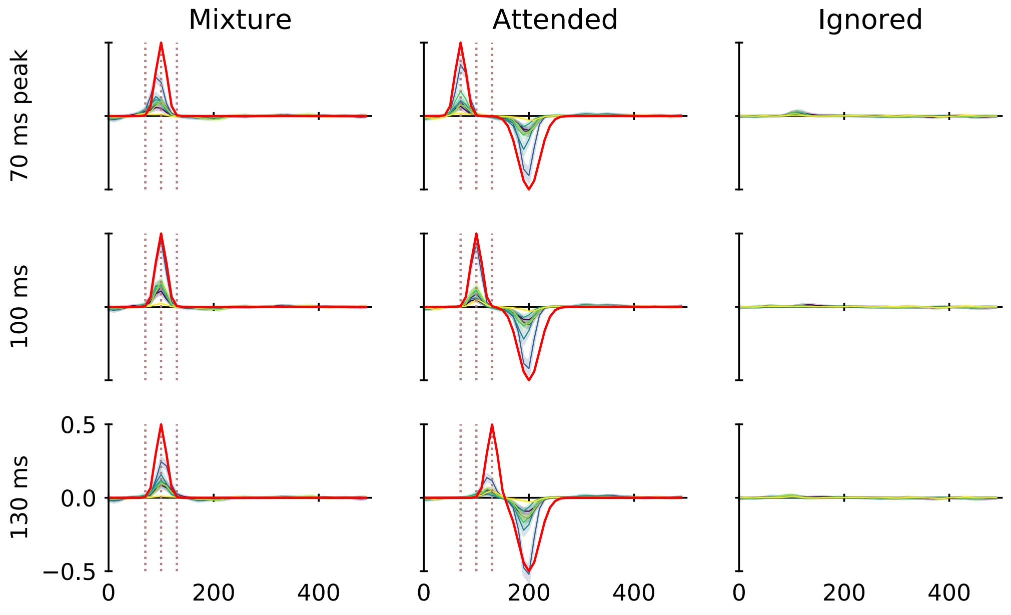


Figure S2. **Simulations with different peak latencies.** Three simulations in which the latency of the attended onsets peak was systematically varied (dotted vertical lines). Simulations were performed to test whether a delay in the response to the attended speech, as observed in the MEG experiment, could be due to how the two responses combine. In each simulation, estimated peak times are slightly biased *towards* the actual peak of the other representation, suggesting that MEG results are not due to such an artifact. Envelopes were included in the simulations as in Figure 1, but responses are not shown here.
